# Formamide significantly enhances the efficiency of chemical adenylation of RNA sequencing ligation adaptors

**DOI:** 10.1101/2022.11.16.516815

**Authors:** Samuel R. Hildebrand, Adam K. Hedger, Jonathan K. Watts, Anastasia Khvorova

## Abstract

Pre-adenylated single-stranded DNA ligation adaptors are essential reagents in many next generation RNA sequencing library preparation protocols. These oligonucleotides can be adenylated enzymatically or chemically. Enzymatic adenylation reactions have high yield but are not amendable to scale up. In chemical adenylation, Adenosine 5□-phosphorimidazolide (ImpA) reacts with 5′ phosphorylated DNA. It is easily scalable but gives poor yields, requiring labor-intensive cleanup steps. Here, we describe an improved chemical adenylation method using 95% formamide as the solvent, which results in the adenylation of oligonucleotides with >90% yield. In standard conditions, with water as the solvent, hydrolysis of the starting material to adenosine monophosphate limits the yields. To our surprise, we find that rather than increasing adenylation yields by decreasing the rate of ImpA hydrolysis, formamide does so by increasing the reaction rate between ImpA and 5′-phosphorylated DNA by ∼10 fold. The method described here enables straightforward preparation of chemically adenylated adapters with higher than 90% yield, simplifying reagent preparation for NGS.

## Introduction

RNA sequencing is a powerful and ubiquitous tool in molecular biology. Next generation sequencing (NGS) libraries are often made by ligating pre-adenylated single-stranded DNA (ssDNA) adapters to RNA 3′ ends. This necessitates either the purchase or synthesis of pre-adenylated ssDNA oligonucleotides (Lau et al. 2001; Viollet et al. 2011; Max et al. 2018). Adenylation can be performed either enzymatically, using various ligases, or chemically, using an activated adenosine 5′-phosphorimidazolide (ImpA) (Lau et al. 2001; Dai et al. 2009; Song et al. 2015; Lama and Ryan 2016; Chen et al. 2012; Max et al. 2018). While the yield of enzymatic adenylation can be high, it is generally performed on a small scale. Chemical adenylation, on the other hand, is scalable but traditionally provides poor yields requiring labor-intensive HPLC or gel purifications post-synthesis (Max et al. 2018).

Here, we report that by performing the chemical adenylation reaction in 95% formamide instead of water, the reaction yield is increased to >90%, removing the necessity of further purification by HPLC or gel extraction. Additionally, we show that the increased yield is due to a faster reaction rate between the 5′ phosphate and ImpA in formamide than in water.

## Results

### Formamide Improves the Yield of Adenylation of 5′-phosphorylated DNA

Chemical adenylation of 5′-phosphorylated ssDNA oligonucleotides in solution is a common method to prepare ligation adapters used in RNA sequencing library preparations. In this reaction, ImpA is reacted with a 5′-phosphorylated ssDNA oligonucleotide to yield a 5′ adenylated product (Figure 1A). This reaction is inefficient and requires laborious purifications to separate unreacted the 5′-phosphorylated oligonucleotide from the adenylated product.

**Figure 1:**
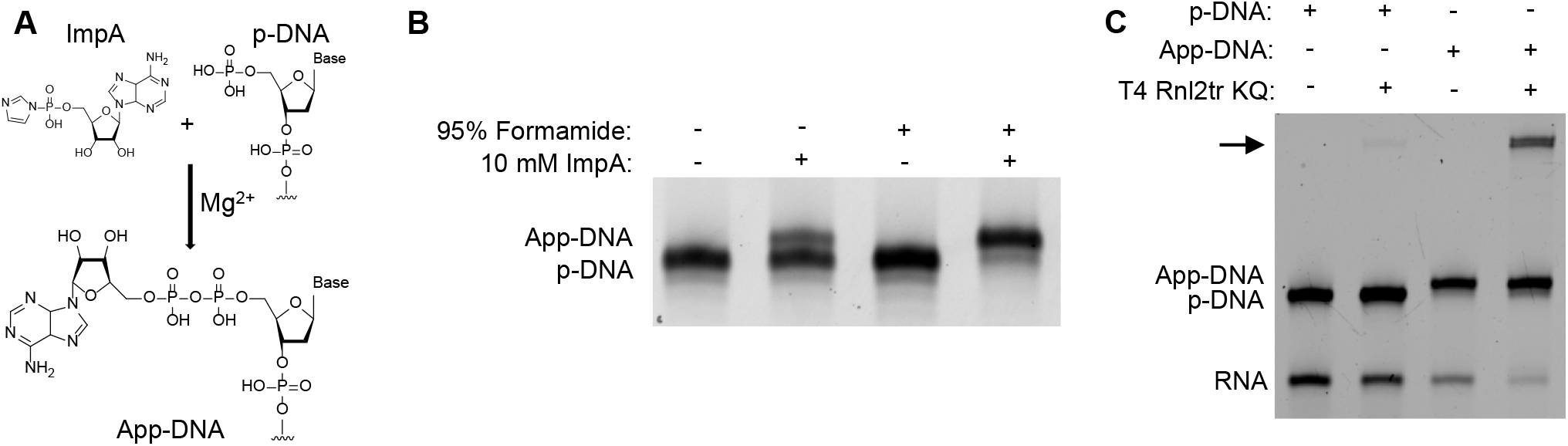
Formamide increases the yield of chemical adenylation. *(A)* Schematic of the adenylation reaction. *(B)* Adenylation of a 5’ phosphorylated DNA oligonucleotide (ODN1) with the reaction taking place either in water or in 95% formamide. *(C)* Ligation of a 5’ p-DNA oligonucleotide or a 5’ App-DNA oligonucleotide that was adenylated in formamide to an RNA oligonucleotide using a mutant RNA ligase that requires a pre-adenylated ligation donor.

Previous studies which focused on adenylating ssDNA oligonucleotides during solid phase synthesis showed some length-dependance on adenylation yield with shorter oligonucleotides being adenylated with higher efficiency. We hypothesized that this length-dependency may be due to increased steric hindrance with longer oligonucleotides and that the low efficiency of the reaction in solution may, similarly, be caused by steric hindrance from oligonucleotide secondary structure. We reasoned that performing the adenylation reaction in denaturing conditions may increase yields due to decreased steric hindrance. To create denaturing conditions, we attempted the reaction in a final concentration of 95% formamide. Analysis by denaturing PAGE showed that the reaction yield was dramatically improved in the presence of formamide as compared to the reaction carried out in water (∼95% vs ∼20%) (Figure 1B). Mass spectrometry confirmed that the product was the adenylated oligonucleotide. To verify that the adenylation occurred on the 5□-position and produced a functional ligation donor, we carried out a ligation reaction using T4 Rnl2tr KQ, a ligase that is dependent on a pre-adenylated donor oligonucleotide. We observed the presence of the proper ligation product in the presence of the pre-adenylated DNA, validating that this method yields a functional adenylated oligonucleotide (Figure 1C).

### The Improved Adenylation Yield of Formamide is Generalizable

To verify that the improvements to the yield of adenylation were generalizable, we tested the adenylation efficiency of a panel of eight 5′-phosphorylated DNA oligonucleotides of varying sequence and length in either 95% formamide or water. Denaturing PAGE analysis showed that formamide drastically increased the adenylation efficiency of all eight oligonucleotides (Figure 2A). The shift seen by denaturing PAGE was verified to be due to adenylation by mass spectrometry (Figure 2B). Interestingly, none of the eight oligonucleotides in the panel had strong secondary structure as determined by *in silico* secondary structure analysis, indicating that the improvement in adenylation yield in formamide may not, in fact, be due to a reduction in secondary structure but rather through another mechanism (Supplementary Figure 1). Intrigued by this, we tested the adenylation reaction with other additives meant to denature oligonucleotide secondary structure and found that while 50% formamide (∼50% yield) and, to a slightly lesser extent, 50% DMSO (∼40% yield) improved adenylation yield, 4M urea did not (∼25% yield) (Supplemental Figure 2).

**Figure 2.**
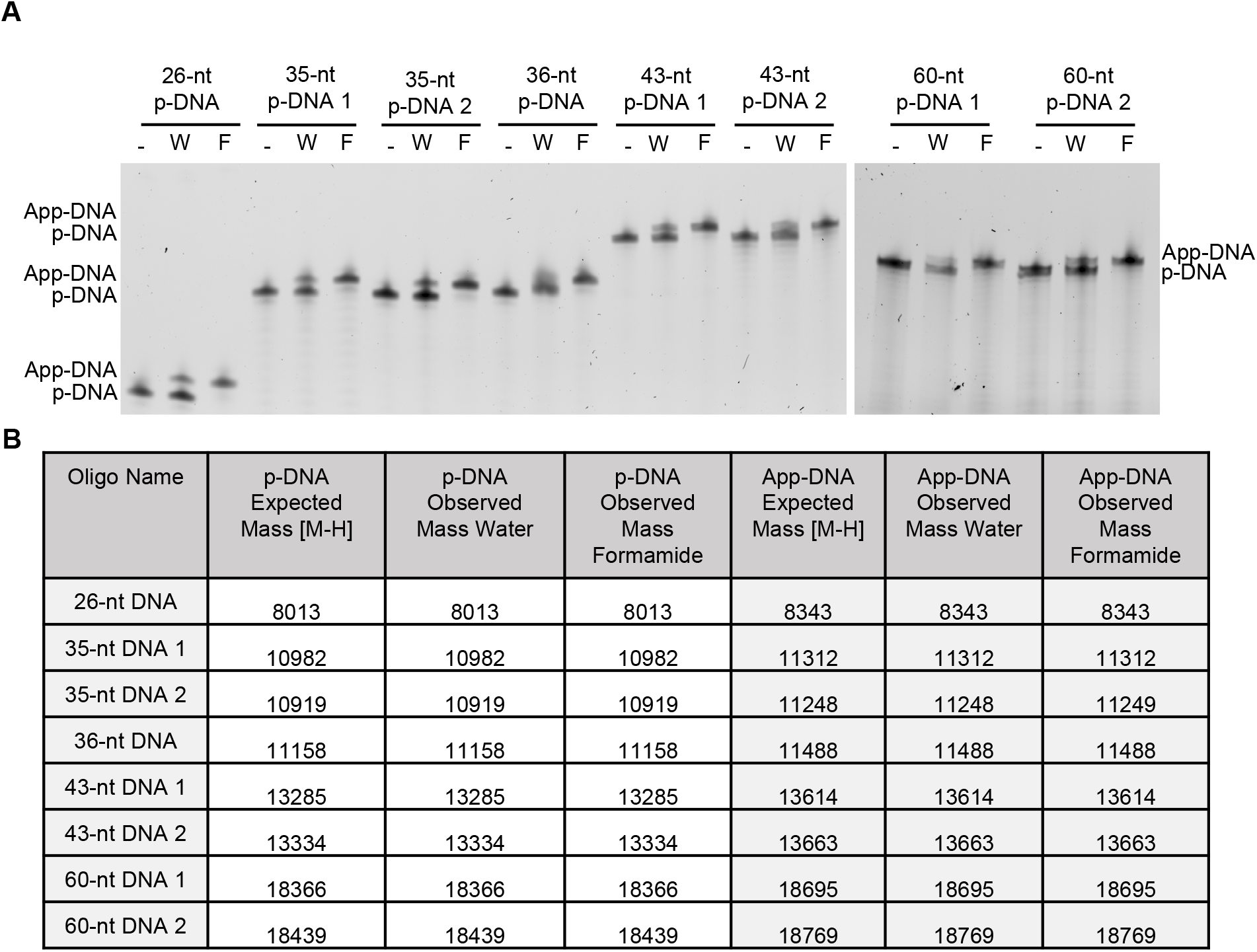
Formamide increases the yield of adenylation across multiple oligonucleotides. *(A)* A panel of 5’ phosphorylated DNA oligonucleotides were adenylated in either water (W) or 95% formamide (F). *(B)* Q-TOF LC/MS results of the adenylation reactions confirms the identity of expected products.

Additionally, when the adenylation reaction was performed in nearly pure formamide (600 mM water from MgCl_2_ hexahydrate solubilized in formamide), a significant fraction (∼20%) of the adenylated species had an additional +28 mass, potentially due to formylation of the oligonucleotide 5□ phosphate or ImpA (Supplemental Figure 2)(Benner et al. 2012). This additional mass is not detected when the reaction is conducted in 95% formamide with 5% water, indicating that the +28 mass is likely hydrolysable. These results indicated to us that the formamide-induced improvement in adenylation yield is due to a mechanism other than secondary structure melting.

### Formamide Increases the Rate of 5′-phosphorylated DNA Adenylation

We reasoned that if formamide is not increasing the yield of the reaction by decreasing secondary structure, it could instead be doing so either by increasing the reaction rate between ImpA and the 5□-phosphorylated DNA or by reducing the rate of degradation of ImpA to adenosine monophosphate (pA). To test if performing the reaction in formamide affects the rate of the reaction between ImpA and 5□-phosphorylated DNA, we performed an adenylation timecourse. The reaction was quenched at various timepoints, resolved by denaturing PAGE, and the adenylation efficiency was measured by densitometry (Figure 3A). Quantification of adenylation showed that the initial reaction rate increases in formamide, with the reaction in formamide proceeding ∼4.25 times faster than the reaction in water (Figure 3B). Additionally, while the reaction in water was fit well by a one-phase exponential, the reaction in formamide was better fit by a two-phase exponential potentially indicating that the adenylation reaction mechanism differs between the two conditions (Figure 3B).

**Figure 3.**
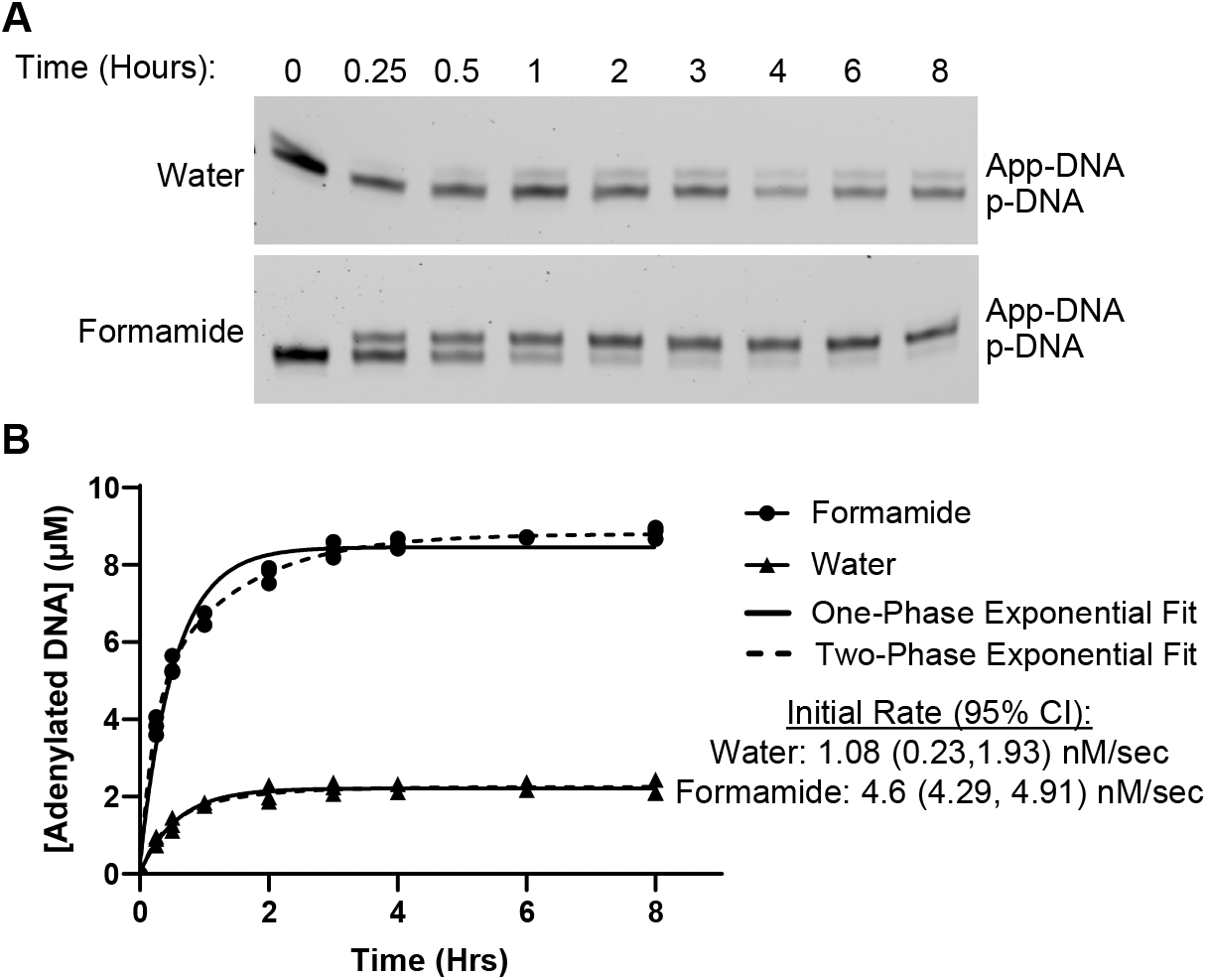
Formamide increases the rate of adenylation. *(A)* Time course of adenylation of a 5′ phosphorylated DNA oligonucleotide (ODN1) in either water or 95% formamide. *(B)* Quantification of the adenylation timecourse showing both one-phase exponential and two-phase exponential equations fit to the data. Mean and confidence intervals for initial rates were calculated by fitting each replicate individually and calculating the initial rate.

### Formamide Increases the Reaction Rate Between ImpA and pA

To gain more insight into the mechanism by which formamide increases the yield of the adenylation reaction, we next chose to track the reaction between mononucleotides over time by HPLC. Since ImpA will react with its hydrolysis product, pA, we are able to measure both the hydrolysis rate of ImpA into pA as well as the reaction rate between ImpA and pA (which acts as a surrogate for 5□-phosphorylated DNA oligonucleotides) to form AppA by using ImpA as the only starting material in the reaction, allowing us to gain insight into the mechanism by which formamide increases the yield of adenylation. From this data and the reaction scheme in (Figure 4), a system of ordinary differential equations (ODEs) was used to fit the reaction rates. The reaction products were resolved by HPLC where clear peaks for ImpA, pA, and AppA can be seen (Figure 5A, Supplemental Figure 3). Plotting the measured data along with the concentrations over time predicted by the ODE fit showed reasonable concordance with the data (Figure 5B). In the water condition, the concentration of AppA is slightly underestimated at early timepoints, while the concentration of pA is slightly overestimated. In the formamide condition, the predicted concentrations better fit the measured concentrations although the level of pA is initially predicted to be too high and is then brought down below the level measured.

**Figure 4.**
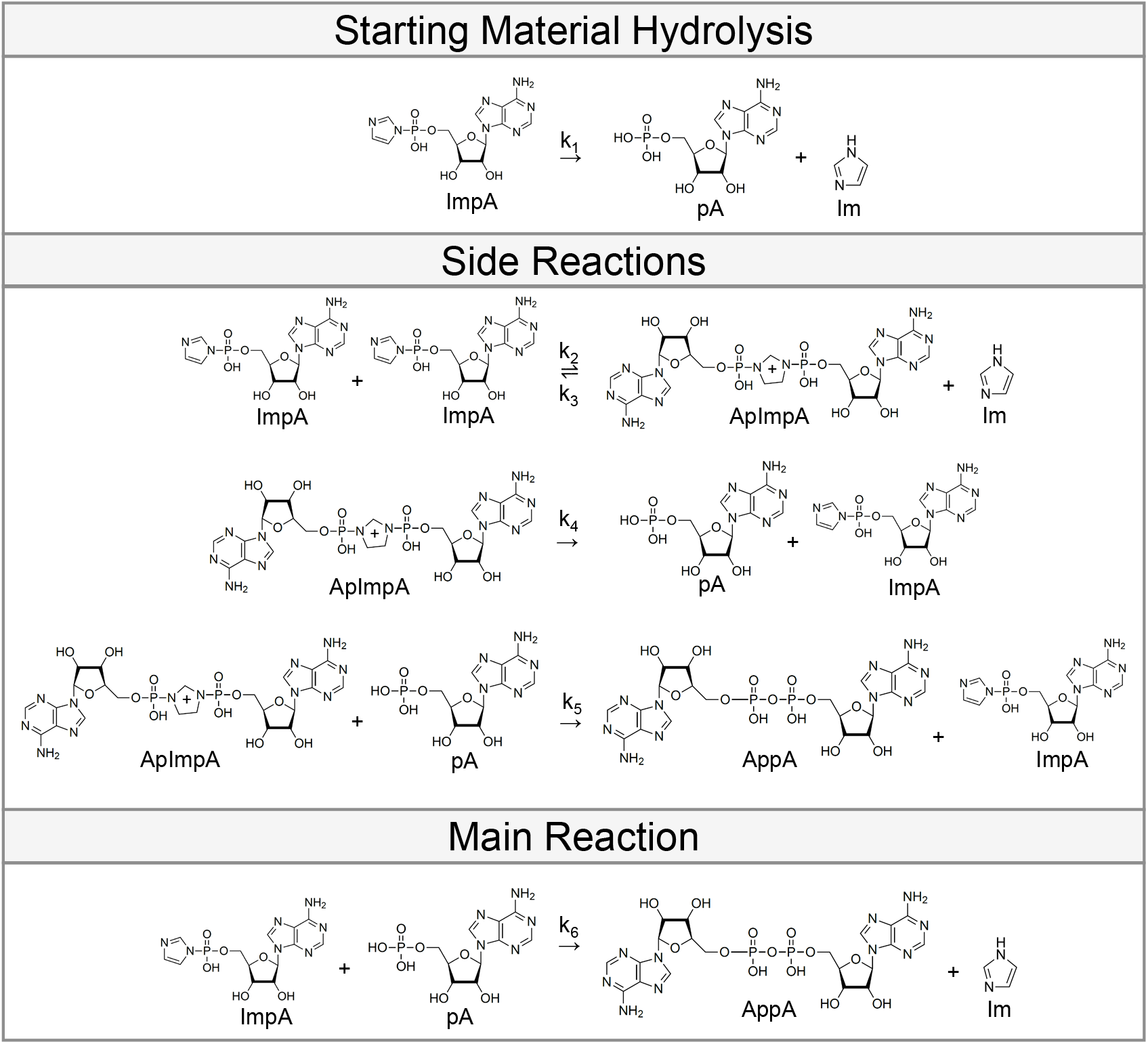
Schematic of the reactions used to fit the system of ordinary differential equations in **(Figure 5)**.

**Figure 5.**
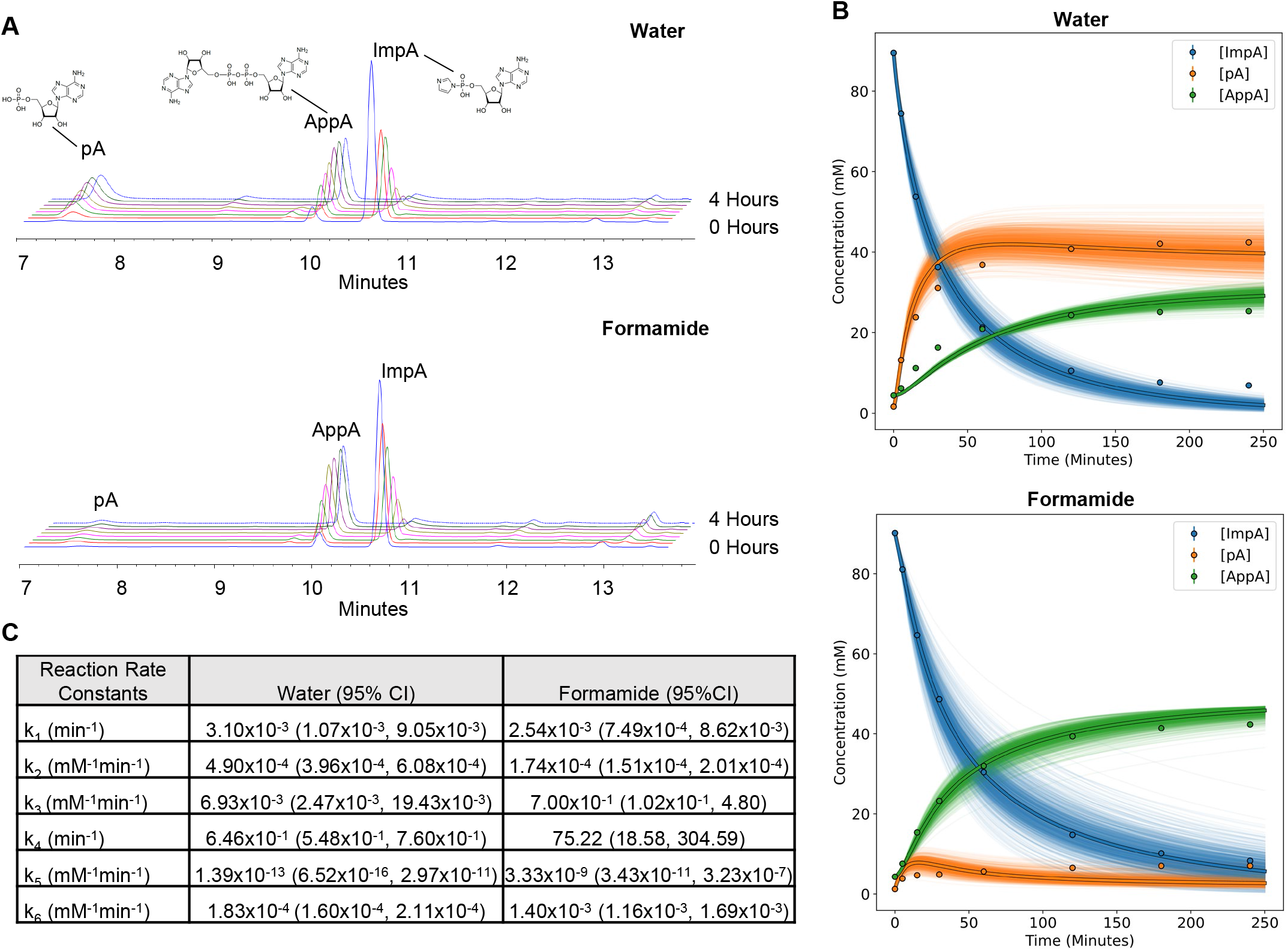
Formamide increases the rate of the reaction between ImpA and pA. *(A)* Representative HPLC profiles of the reaction between ImpA and its degradation product, pA, to make AppA in either water or 95% formamide over a 4 hour timecourse. *(B)* Concentrations of ImpA, pA, and AppA predicted (lines) by fitting the reaction scheme in **(Figure 4)** to the HPLC data (points) with a system of ordinary differential equations. The bold lines are the concentrations predicted by the mean of the log-transformed fit rates, and the faint lines are the predicted concentrations from resampling the rates from their 95% confidence intervals. *(C)* Reaction rates for the scheme in **(Figure 4)** fit by the system of ordinary differential equations.

The reaction rates calculated from the ODE fit showed that the fitted rate of ImpA decay was no different in formamide and water while the predicted reaction rate between ImpA and pA was nearly ten times faster in formamide than in water (Figure 5C). This data agrees well with the oligonucleotide adenylation timecourse data in (Figure 3) and provides evidence that the main driver in the increased adenylation yield is a higher reaction rate between ImpA and 5□ phosphorylated DNA.

## Discussion

Chemical adenylation of DNA oligonucleotides using ImpA has long been thought of as an inefficient reaction requiring additional purification to remove unadenylated oligonucleotides. Even though the reaction on solid support has been optimized, it still yields <50% adenylated product for long DNA oligonucleotides, such as those used as adapters in RNA sequencing library preparations (Dai et al. 2009). In this work we show that by performing the adenylation reaction in 95% formamide, the yield is improved to >90% regardless of oligonucleotide length and sequence. This improvement will make the production of 5′-adenylated oligonucleotides, a key reagent in NGS, easier and more accessible.

After verifying the generalizability of formamide for improving adenylation yields we explored the mechanism of this improvement. The adenylation efficiency could not be explained by a reduction in secondary structure since the efficiency is improved in unstructured DNA oligonucleotides. Additionally, other agents that disrupt secondary structure do not provide the same degree of improvement as formamide. This suggested that formamide acts either to improve the reactivity between the 5′-phosphate and ImpA or the stability or ImpA, allowing more to react before it is hydrolyzed. Kinetic experiments revealed that the improved adenylation efficiency is driven by an increased reaction rate between the 5′-phosphate and ImpA in formamide as compared to water, not a change in the hydrolysis rate of ImpA.

Mass spectrometry results from oligonucleotides adenylated in pure formamide show a +28 mass, which is not detectable when performing the reaction in 95% formamide, likely corresponding to formylation of the adenylated product. Reactions between formamide and nucleotides have long been studied in the context of prebiotic chemistry, including the finding that it is a suitable solvent for the phosphorylation of nucleosides in the presence of inorganic phosphate (Xu et al. 2019; Schoffstall 1976). While it remains unclear whether formamide is increasing the reaction rate by formylating either the oligonucleotide or ImpA or through a solvent effect, the presence of this hydrolysable +28 adduct raises the possibility that formamide is participating in the reaction as an activator. In keeping with the idea of formamide acting as an activator to increase adenylation yields, recent studies have also shown that the addition of high concentrations of 1-methyl-imidazole as an activator increases the yield of an RNA capping reaction between 7-methylguanosine 5′-diphosphate imidazolide and a 5□-phosphate RNA (Abe et al. 2022). Formamide can, however, formylate nucleotides at multiple sites, making it difficult to assess what the identity and role, if any, of this formylation may be (Benner et al. 2012).

This improved method allows for the large scale and facile synthesis of 5′-adenylated oligonucleotides, making it easier to synthesize large amounts of many adenylated oligonucleotides simultaneously without the additional step of HPLC or gel purification. This will allow researchers to spend less time and effort on reagent preparation when making RNA sequencing libraries.

## Materials and Methods

### Oligonucleotides

All oligonucleotides used in this study were purchased from Integrated DNA Technologies (IDT) as 5□-phosphorylated DNA with standard desalting.

*Synthesis of ImpA*. Adenosine 5′-monophosphoric acid (0.5 mmol, 174 mg, 1.0 eq) in 15 mL DMF was added dropwise to a vigorously stirred solution of triphenylphosphine (1 mmol, 262 mg, 2.0 eq), 2,2′-dipyridyldisulfide (1 mmol, 220 mg, 2.0 eq), and imidazole (2.5 mmol, 170 mg, 5.0 eq) in 15 mL DMF and triethylamine (2.5 mmol, 0.9 mL, 5.0 eq). This solution was then stirred for 1.5 hours at room temperature. ImpA was then precipitated adding the reaction to a stirred solution of 53 mM sodium perchlorate in 115 mL acetone and 55 mL anhydrous ethyl ether in a dropwise manner. The precipitate was then collected by passing it through a Büchner funnel and washed thoroughly with acetone. Finally, the precipitate was washed with anhydrous ethyl ether and dried under a vacuum overnight. Purity of the final product was assessed by phosphorous NMR and proton NMR (Tabulation below, full spectra in Supplemental Figure 4). ImpA was then stored under argon at -20°C.

^**1**^**H NMR** (DMSO-*d*_6_, 500 MHz) δ_ppm_ 8.40 (1H, s), 8.14 (1H, s), 7.63 (1H, s), 7.29 (2H, brs), 7.09 (1H, brd, *J* = 0.8 Hz), 6.85 (1H, s) 5.89 (1H, d, *J* = 6.2 Hz) 5.48 (1H, d, *J* = 6.2 Hz), 5.30 (1H, d, *J* = 4.4 Hz), 4.58 (1H, dd, *J* = 5.8 Hz, 5.3 Hz), 4.03 (1H, d, *J* = 3.2 Hz), 3.94 (1H, d, *J* = 3.2 Hz) 3.73 (2H, m)

^**31**^**P NMR** (DMSO-*d*_6_, 202 MHz) δ_ppm_ -10.35 (1P, s)

### Oligonucleotide Adenylation Reaction

Adenylation reactions were carried out by mixing 100 mM MgCl_2_•6H_2_O (from a 1M solution made either in water or formamide), 10 mM ImpA (from a 100 mM solution made either in water or deionized formamide (Invitrogen, AM9344)), and 10 µM 5′-phosphorylated DNA (from a 200 µM solution made in water) either in water or deionized formamide. This solution was then heated to 65°C for the indicated times. The reaction was then diluted 1:3 in water, ethanol precipitated, and resuspended in 95% formamide loading dye. The adenylation reactions were resolved by 7.5M urea 15% PAGE and stained with Sybr Gold (Thermo S11494).

### T4 Rnl2KQ Ligation

Ligations were performed by mixing 1 µL 10 µM of DNA adapter, 1 µL 10 µM RNA oligonucleotide, 3 µL 10x RNA ligation buffer (500 mM Tris-HCl pH 7.5, 100 mM MgCl_2_, 100 mM DTT), 15 µL 50% PEG 8000, 3 µL Rnl2KQ (NEB M0373S), water to 30 µL. The ligations were incubated for 1 hour at room temperature and quenched by boiling with 30 µL 100% formamide and 1 µL 10 mM EDTA. The reactions were resolved by 7.5M urea 15% PAGE and stained with Sybr Gold (Thermo S11494).

### Nucleotide Adenylation Reaction and HPLC

Reactions were carried out by mixing 1M MgCl_2_, 100 mM ImpA in either water or 95% deionized formamide. The solution was heated to 65°C for the indicated times. Timepoints were quenched by adding 10 µL of the adenylation reaction to 490 µL 50 mM EDTA and flash freezing in liquid nitrogen. HPLC was performed by individually thawing the samples and analyzing them immediately. Samples were injected on a 1260 HPLC (Agilent) using a Luna Omega PS-C18 5 µm 100 Å liquid chromatography column (250 × 4.6 mm) (Phenomenex). Samples were resolved over 18 minutes using an isocratic elution gradient of 2% for 3 minutes, followed by 2-20% over 15 minutes (95% acetonitrile, 5% 15 mM ammonium acetate pH 4.2) in 15 mM ammonium acetate pH 4.2 at 25°C, using a flow rate of 1.0 mL min^-1^. This was followed by wash steps at higher elution strengths between runs. Absorbance was recorded at 260 nm and peaks were collected manually for LCMS analysis. The order of elution was pA (∼7 minutes), followed by AppA (∼9.5 minutes), and lastly ImpA(∼10.5 minutes).

For High-resolution mass spectrometry (HRMS) analysis, nucleotides were first resolved using the HPLC method above, and UV peaks were manually collected in separate vials. These samples were immediately run on a 6530 Accurate-Mass Q-TOF LC/MS (Agilent) on positive ESI mode linted to a pre-injection 1260 infinity HPLC (Agilent). Mass spectrometry settings to detect nucleotides were as follows: gas temperature 350°C, nebulizer gas rate 45 psig, drying gas 10 L/min, VCap 4000 V, fragmentor voltage 120 V, skimmer voltage 65 V.

### Ordinary Differential Equation Fitting for Nucleotide Adenylation Reaction Rates

To calculate the reaction rates for the nucleotide adenylation reaction, the following system of ordinary differential equations were fit to the HPLC data using the ODE45 function in Matlab R2021b:

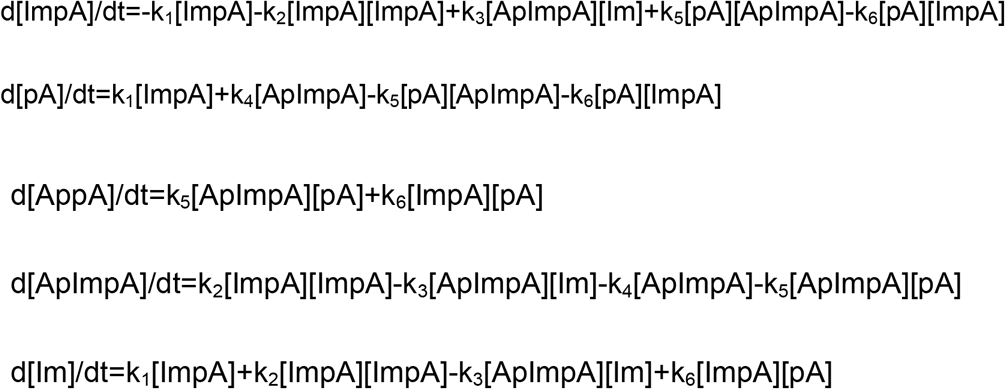

Initial conditions were set as follows:

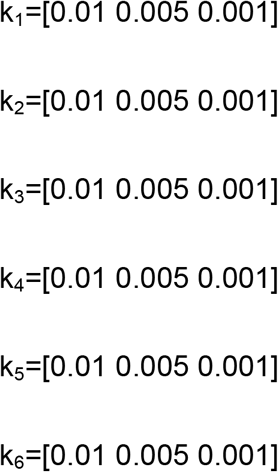

All combinations of initial rates were used to initialize the ODE45 optimization and the median of all fits that reached a local minimum or were stopped due to the step size being below 1×10^−10^ was calculated for each rate constant in each replicate.

The log(mean) of the fitted rates were then used to calculate confidence intervals and 1000 new rates were sampled from the 95% confidence interval for each rate constant. The mean rates as well as the resampled rates from the 95% confidence interval were used to plot the predicted reactant concentrations against the measured concentrations using the Scipy and Matplotlib packages in Python 3.7.

### LC–MS analysis of oligonucleotides

The mass of oligonucleotides was measured by LC–MS analysis on an Agilent 6530 accurate mass Q-TOF using the following conditions: buffer A: 100□mM 1,1,1,3,3,3-hexafluoroisopropanol (HFIP) and 9□mM triethylamine (TEA) in LC–MS grade water; buffer B:100□mM HFIP and 9□mM TEA in LC–MS grade methanol; column, Agilent AdvanceBio oligonucleotides C18; linear gradient 0–100% B 8min; temperature, 60°C; flow rate, 0.5□ml/min. LC peaks were monitored at 260nm. MS parameters: Source, electrospray ionization; ion polarity, negative mode; range, 100–3,200□m/z; scan rate, 2 spectra/s; capillary voltage, 4,000; fragmentor, 180□V.

## Acknowledgements

We wish to thank Dr. Li Li for his invaluable advice throughout the project. This work was supported by funding from the National Institutes of Health (NIH) grants R35-GM131839 (A.K.), S10-OD020012 (A.K.), F31-NS122493 (S.R.H.), R01-NS111990 (J.K.W.).

---

**Supplemental Figure 1:**
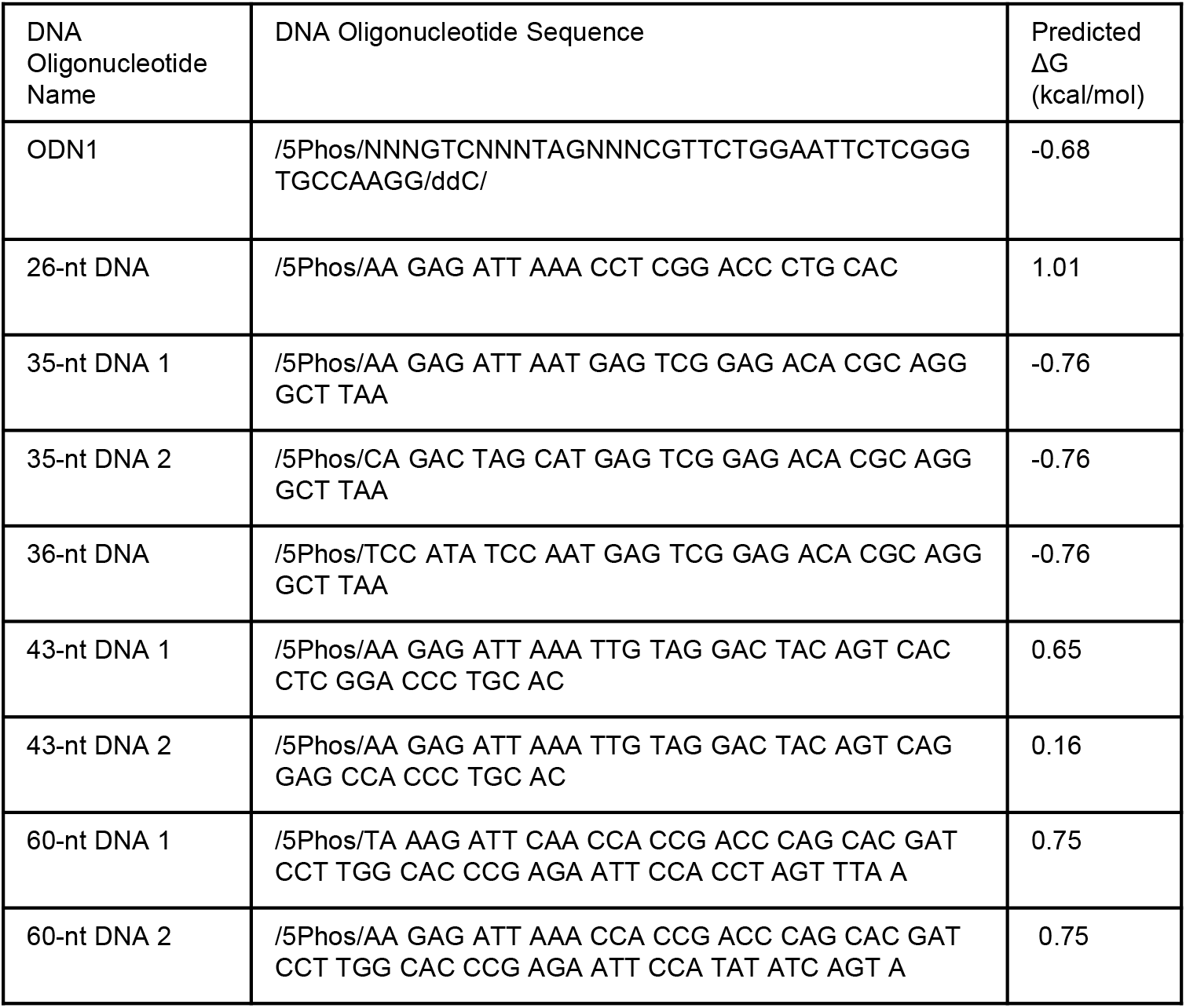
Sequence and predicted ΔG of the oligonucleotides used in the adenylation reactions.

**Supplemental Figure 2:**
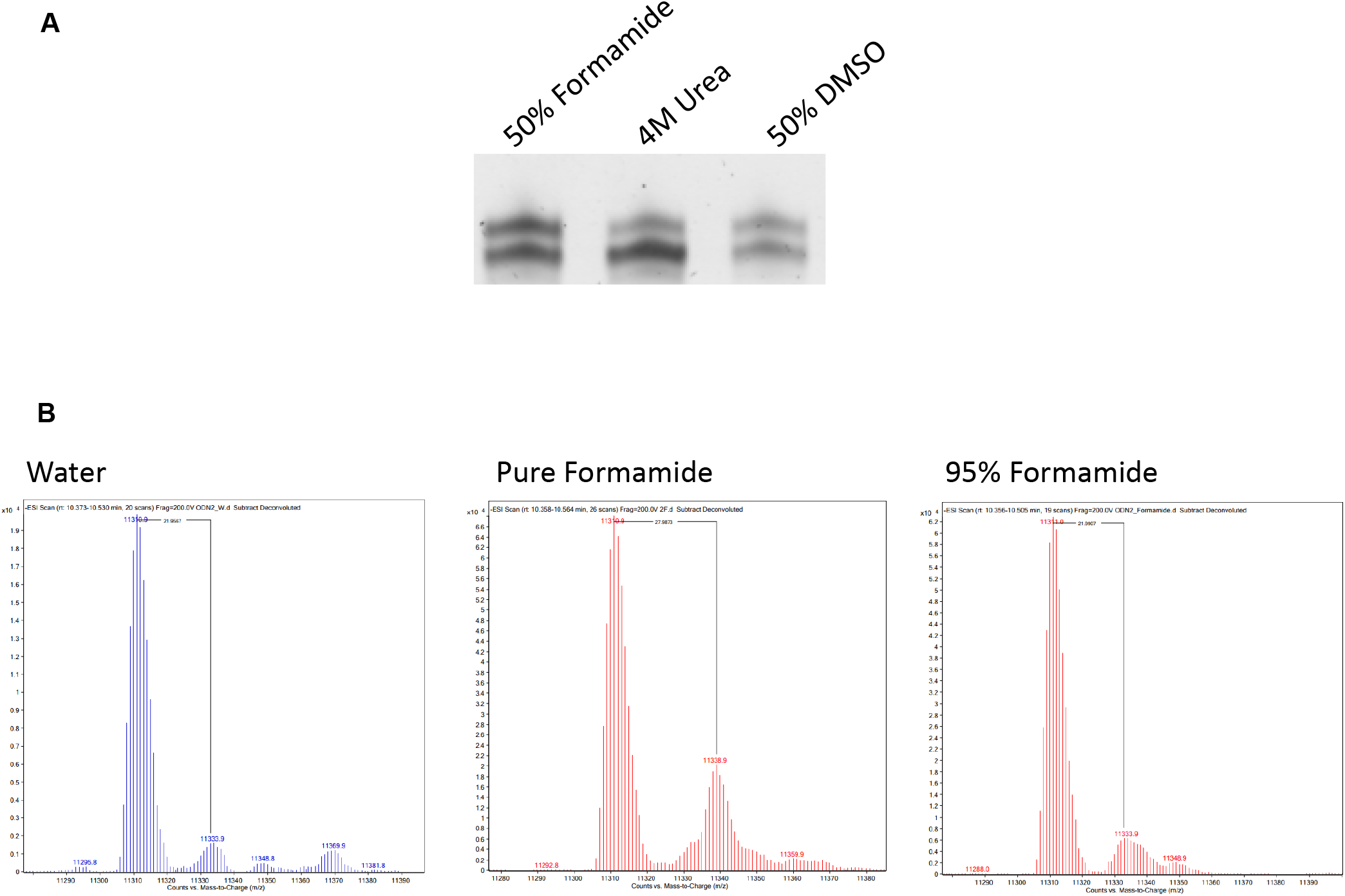
*(A)* Adenylation of a 5’ phosphorylated DNA oligonucleotide in the indicated solvents. *(B)* Q-TOF LC/MS results of a 5’ phosphorylated DNA oligonucleotide (35-nt DNA 1) adenylated either in water, pure formamide, or 95% formamide showing the presence of a mass of the adenylated product +28 when the reaction is performed in pure formamide.

**Supplemental Figure 3:**
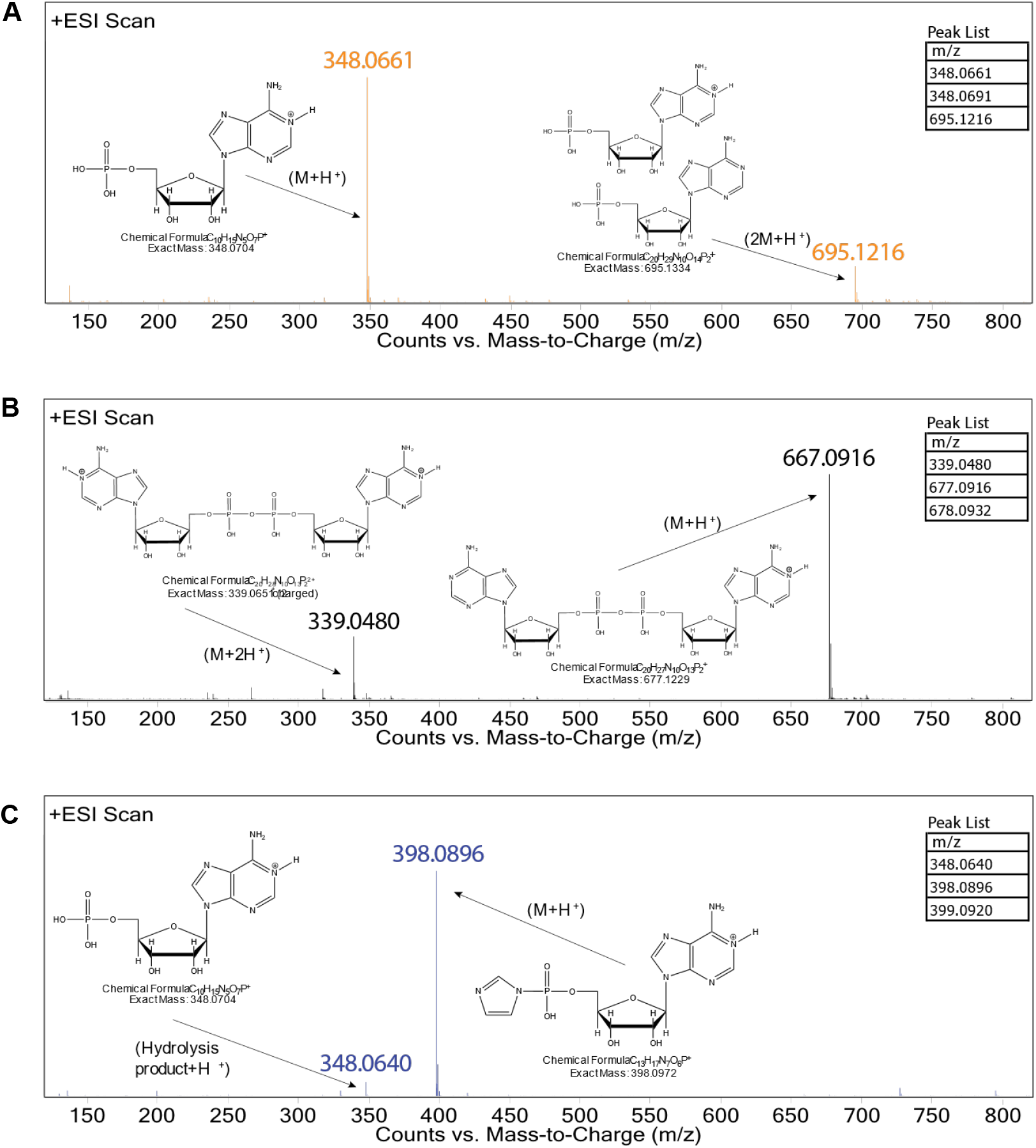
High-resolution Mass Spectrometry to identify the HPLC peaks in **(Figure 5)**. *(A)* Results from the ∼7-minute HPLC peak. *(B)* Results from the ∼9.5-minute HPLC peak. *(C)* Results from the ∼10.5-minute HPLC peak.

**Supplemental Figure 4:**
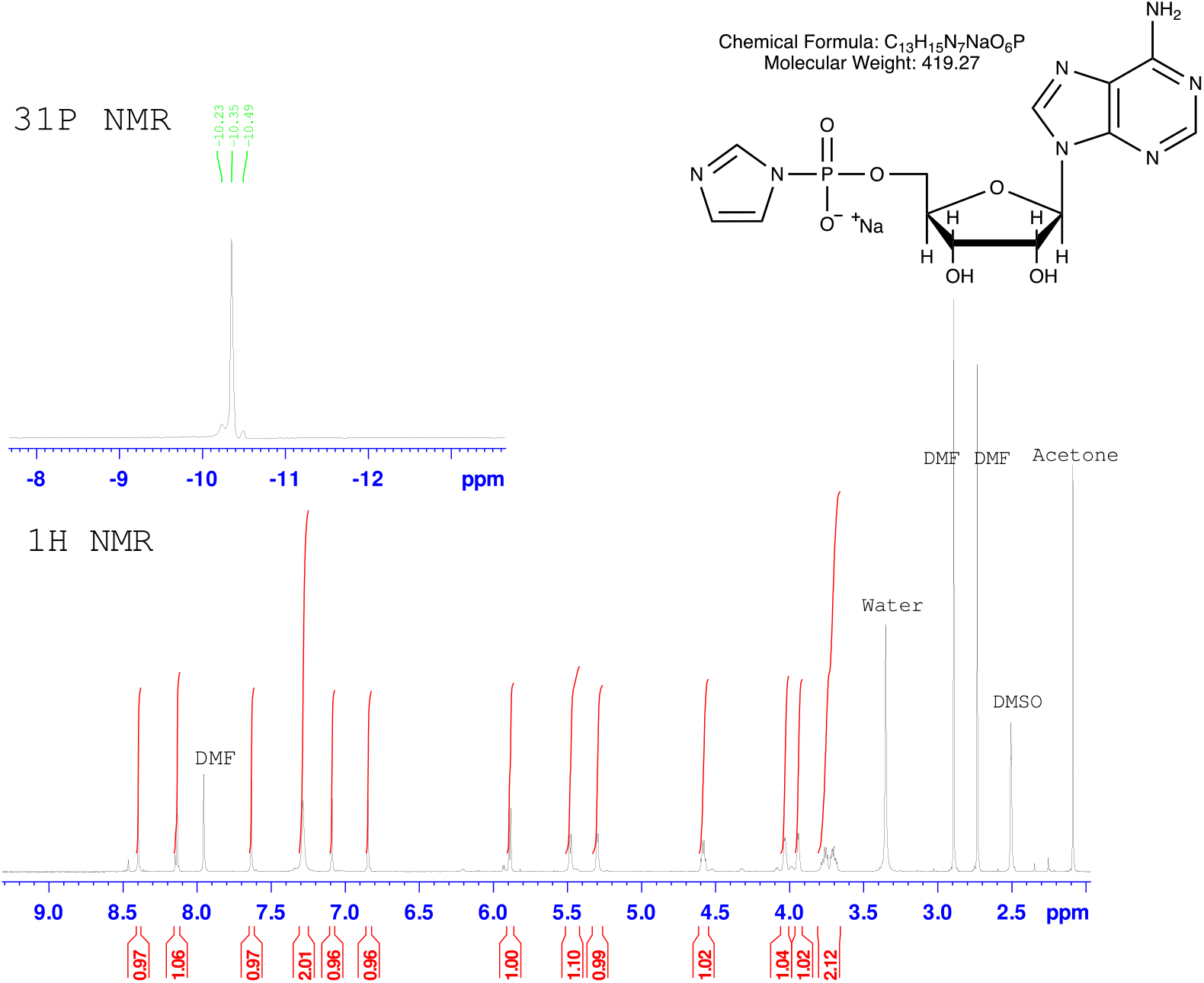
^1^H and ^31^P NMR of adenosine 5’-phosphorimidazolide (ImpA) in DMSO-*d*_*6*_.

